# In situ vaccine effect of radiotherapy is associated with intratumoural ERV transcription and RNA virus sensing

**DOI:** 10.64898/2026.02.06.704391

**Authors:** Stella Logotheti, Elif Yildiz, Sarah Hasan, Elpida Theodoridou, Stefan Kuhn, Thorsten Stiewe, Stephan Marquardt, Athanasia Pavlopoulou, Joao Seco

## Abstract

Radiotherapy (RT) transforms tumour tissues into *in situ* vaccines that trigger antitumor immunity. Immunogenicity depends on how RT is delivered, since heterogeneous RT (spatially fractionated RT, SFRT) elicits more prominent responses than the conventional homogenous one. However, this phenomenon cannot be clinically harnessed, unless the relevant pathways are identified. To gain insights, we developed a hybrid dry-lab/wet-lab approach that integrates systems-level immune phenotypes established by homogenous or heterogenous RT (SFRT) with the transcriptomic profiling of irradiated tumors. By further combining feature extraction with machine-learning, including multilayer perceptron modelling, we ranked predictors of immune infiltration and patient survivability for each RT type. We found that conventional RT induces coordinated upregulation of cytosolic sensors of RNA viruses (OASes and RIG I-like receptors) along with ERV RNAs predominately 400-800 base-pairs long, which might serve as their ligands. For schemes establishing abscopal effects, a coordinated upregulation of the OAS sensors and shared ERV transcripts was observed in both irradiated and distant tumours. Compared to homogenous RT, SFRT triggered earlier and stronger activation of OAS signaling along with NK cell responses. Overall, we show a co-involvement of tumour cell-intrinsic ERVs and their cytosolic RNA sensors in RT-induced antitumor immunity. This key finding could guide mechanistic studies and future precision oncology.

## INTRODUCTION

Radiotherapy (RT) is traditionally employed in the clinical setting for cancer treatment either as a standalone practice or in combination with other therapeutic modalities. Beyond its conventional cytotoxic role, RT reprograms tumours into immunogenic tissues, which train the immune system to recognize and eradicate cancer cells within and beyond the irradiated site. At rare clinical cases, tumours outside the irradiated area respond to RT, a phenomenon known as “abscopal effect” ^1^. Hence, RT has re-surfaced as a powerful means to enhance the efficacy of immunotherapeutics, because its combination with immune checkpoint inhibitors is envisaged to overcome resistance to immunotherapy and increase the frequency of abscopal effects ^1^. The signaling pathways through which RT transforms tumours to *in situ* vaccines capable or eliciting durable antitumour immunity have remarkable translational value, and their elucidation is essential to guide the design of personalized radioimmunotherapy regimens.

The immune response to RT entails tumour cell-intrinsic reprogramming, cell-cell interactions in the tumour microenvironment (TME), and systems-level regulation of the immune system. Induction of the DNA damage response and repair (DDR/R) in the irradiated tumour is fundamental ^2^, as it triggers immunogenic cell stress and death (ICD) ^3^. RT-activated DDR/R promotes cell cycle arrest and death. In turn, stressed and dying cancer cells produce and release a repertoire of bioactive molecules, including but not limited to Damage-Associated Molecular Patterns (DAMPs), that act on immune cells to elicit antitumour responses ^3^. Once the innate immune system senses DAMPs through sensors, called pattern recognition receptors (PRRs) in a motif-specific manner, it promotes a pro-inflammatory milieu and sterile inflammation. DAMPs further activate cytotoxic T lymphocytes, promoting immunological memory and adaptive immune responses ^3^. Transcriptional reprogramming of the irradiated tumour is a key event, as the transcriptomic profiles induced by different RT types and doses trigger divergent signalling cascades, ultimately establishing differential tumour-immune cell interactions and therapeutic outcomes ^4^.

A central signal transduction pathway linking RT-induced DNA damage with immune responses via transcriptional reprogramming is the interferon (IFN) cascade. Cytoplasmic DNA fragments are sensed by the cyclic GMP-AMP synthase (cGAS) which activates the stimulator of interferon genes (STING). STING subsequentlyrecruits kinases like TBK1 (TANK-binding kinase 1), which in turn phosphorylate the transcription factor IRF3 (interferon responsive factor 3). The activated IRF3 drives the transcription of a set of immunomodulatory genes, leading to interferon secretion and immune response ^5^. In certain cancer types, RT also depends on RNA sensors, whereby ionizing radiation amplifies type I interferon signalling via the RIG-I/MAVs-dependent RNA-sensing pathway ^6^. Most interestingly, transposable elements (TEs), particularly transcripts from long-terminal repeats (LTRs) serve as the RNA ligands of RIG-I that trigger this cascade ^6^. TEs reside in the so-called “junk DNA” regions and are emerging as under-noticed instructors of the immune system ^7^. Although most TE copies are non-functional, a subset can be mobilized and transcribed in cancer cells. RT-mediated activation of these elements generates a reservoir of immunogenic RNA species capable of eliciting IFN cascades via RNA-protein interactions ^7^. This novel non-coding RNA arm of RT-induced immunogenicity remains largely enigmatic, compared to the well-characterized cGAS/STING pathway.

Tumour immunogenicity depends on the delivery mode of RT, whether homogenous or heterogeneous ^8^. Spatially heterogeneous distribution of RT (spatially fractionated RT, SFRT) has fewer side effects and elicits stronger anti-tumor responses that the traditional, homogenous RT. In colorectal cancer mouse models, simultaneous administration of high-dose to one half of the tumour and low-dose to the other half, generated a more reactive tumour immune microenvironment compared to the homogenously irradiated controls ^8^. Other studies used microbeam (MRT) or minibeam (MBRT), where a collimator delivers extremely high-dose-rates into high-dose areas (‘peaks’), separated from low-dose regions (‘valleys’) by a few hundred micrometers or millimiters, correspondingly ^9^. In preclinical rodent models, this type of spatial fractionation consistently induced superior *in vivo* antitumour immunity phenotypes ^10-12^. Most recently, minibeam RT achieved a complete clinical response and triggered abscopal effects in a patient with recurrent metastatic sinosinal melanoma who had been unresponsive to conventional X-rays combined with checkpoint inhibitors (CPIs) ^13^. These findings highlight a novel paradigm, in which heterogeneous intratumour irradiation represents an appealing immunotherapy partner, due to its reduced toxicity and increased immunogenicity ^14^.

Uncovering the transcriptional pathways activated in tumours that drive *in situ* vaccine effects, in relation to irradiation type, dose and delivery mode, is urgently needed. Herein, we leverage studies on murine cancer models in which homogeneous or heterogeneous RT elicited broad antitumour immunity throughout the organism. We associate the phenotypes of RT-induced anti-tumour immunity and abscopal effects with transcriptional changes in both protein coding genes and TEs within the targeted tumours. Our analysis unveiled in detail the transcriptional reprogramming induced by conventional X-rays and SFRT and highlighted key differences in terms of immunogenic response to RT and survivability prediction in patients.

## RESULTS

### Broad antitumour immunity phenotypes are associated with tumour transcriptional reprograming in response to conventional RT and SFRT

We mined for all the publicly available studies where rodent tumour models received any type of RT and demonstrated robust phenotypes of local and/or systemic anti-tumour immune responses. Eight studies across six cancer types with their corresponding bulk transcriptomics data (RNS-Seq or RNA microarrays) were eligible ^10,12,15-20^ (Table 1), whereby conventional X-ray irradiation (CONV) was applied in a total dose range of 8-30 Gy. In the lymphoma study ^19^, one sequenced tumour was located within the RT-targeted field and the other one outside the field. This setting allowed us to detect DEGs in non-irradiated (NIR) tumours that represent reporters of abscopal effects. In two studies ( ^10^ and GSE56113), CONV was compared to microbeam RT (hereafter referred to as SFRT), providing insights into transcriptional reprogramming associated with the in situ vaccine effects of each dose delivery modality. For each dataset, transcriptomic analyses identified up- and down-regulated transcripts (logFC < -1.5 and > 1.5; p value < 0.05) in treated versus untreated controls. The number of DEGs varied across cancer types, perhaps due to differences in cancer type, irradiation dose, and transcriptomic platforms.

**Table 1.** Overview of studies where mouse phenotypes of in situ vaccine effects have publicly available transcriptomics data of the ‘immunogenic’ targeted tumours.

| Study accession number | Cancer type | Model | Irradiation Type | Transcriptomics platform | Time of sample collection post-RT | Sample undergoing RNAseq (irradiated vs non-irradiated controls) |
| --- | --- | --- | --- | --- | --- | --- |
| GSE207717 | Pancreatic adenocarcinoma | KPC syngeneic mouse (strain C57BL/6J) and PDAC organoids derived from KPC-OG mice (KPOG) | X-rays, 1 fraction of 8Gy | Illumina NovaSeq 6000 (Mus musculus) | 24 and 48 h post irradiation | KPOG organoids |
| GSE281695 | Lymphoma | BALB/c mice with two A20 lymphoma tumours | X-rays, 1 fraction of 8Gy or 2 fractions of 4 Gy | Illumina NovaSeq X Plus (Mus musculus) | 24 hours and 7 days post treatment. | Tumour inside and outside the irradiated area |
| GSE168016 | Colorectal | MC38 syngeneic mice (strain C57BL/6J) | X-rays, 1 fraction of 15 Gy | Illumina NovaSeq 6000 (Mus musculus) | 60 hours | tumour cells |
| GSE209765 | Colon | CT26 syngeneic mice (BALB/c or C57BL/6) | X rays, 1 fraction of 8Gy | DNBSEQ-G400 (Mus musculus) | When tumour reached 2000 mm<sup>3</sup> | Tumour treated with mock immunotherapy and received RT two days later |
| GSE268311 | Melanoma | B16 melanoma mice (strain C57BL/6) | SFRT (microbeams) vs X-rays | NanoString nCounter® XT Assay | 2 days and 7 days post-RT. Non-irradiated controls collected 12 days post-tumour implantation. | Tumours |
| GSE56113 | Glioblastoma | orthotopic rats bearing intracranial 9L brain tumours | SFRT (microbeams) vs X-rays | Affymetrix GeneChip® Rat 230_2 | 6 hours, 48 hours, 8 days and 15 days after treatment | tumours |
| GSE61208 | Breast | 4T1 syngeneic mice | X-rays, 5 x 6 Gy | [Mouse430_2] Affymetrix Mouse Genome 430 2.0 Array | 4 days post-RT | Tumours |
| GSE255242 | Breast | Murine transplantable TS/A tumour-cell-line co-engrafted with CAFs in syngeneic mice | X-rays, 2 x 6 Gy | NextSeq 2000 (Mus musculus) | when the tumour size reached 2000 mm<sup>3</sup>. | Tumour treated with RT when reached a volume of 80-100 mm<sup>3</sup> |

To extrapolate the transcriptional changes related to the mouse anti-tumour immunity phenotypes to humans, we identified the human orthologues of the RT-upregulated transcripts and performed downstream functional in silico analyses according to the pipeline described in Fig. 1A. Most upregulated rodent genes have human counterparts, while the small percentage of non-conserved genes may account for rodent-specific responses (Fig. 1B). Interestingly, although CONV transactivates relatively few genes in melanoma and glioblastoma, this trend is reversed by SFRT, implying the ability of the latter to boost transcriptional activation (Fig. 1C). We then pooled the human homologues of all CONV-upregulated (across six cancer types) and the SFRT-upregulated transcripts (across two cancer types). We found a substantial overlap of genes transactivated by both RT types (Fig. 1D), which are associated with the activation and/or regulation of several immune cells, such as lymphocytes, T cells and mononuclear cells (Fig. 1E). There were also genes specific for each RT modality. Hence, based on the above observations, we postulated that some of these genes may account for the reported differential effects of SFRT versus CONV and proceeded to analyse the transcriptional responses for each modality separately.

**Figure 1:**
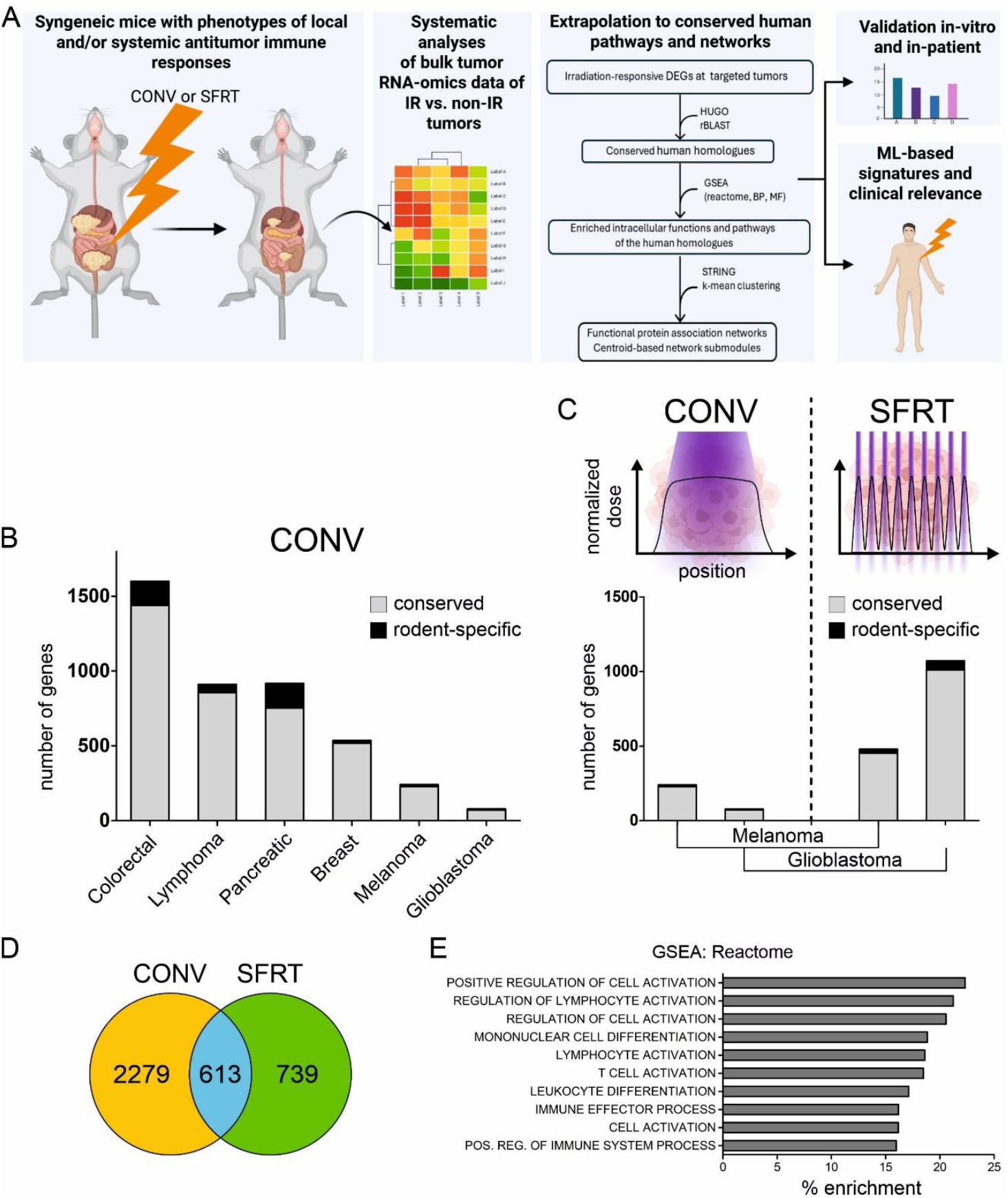
Association of antitumour immunity/abscopal phenotypes with transcriptional changes in response to CONV RT vs. SFRT. (A) workflow of the systemic meta-analysis (created with Biorender). RNAseq data from irradiated tumors extracted from mice with phenotypes of antitumor immune response were compared to their non-irradiated controls. The human orthologues of upregulated genes were defined and subjected to functional enrichment and network analysis, as well as machine learning-facilitated correlations with clinical patient data, (B) number of CONV RT-upregulated transcripts per analyzed cancer type. (C) number of SFRT-vs. CONV RT-upregulated transcripts in glioblastoma and melanoma. The grey columns represent rodent genes with conserved human orthologues (D) Venn diagram of conserved genes upregulated by SFRT vs. CONV RT (E) GSEA reactome analysis of the 613 genes shared between CONV and SFRT

### Conventional radiotherapy activates tumour cell-intrinsic pathways of IFN-dependent RNA viral immune responses

Analysis of CONV-treated tumours versus their untreated contols across the six types of cancers (Table 1) identified 161 genes that were commonly upregulated in ≥ 3 of the 6 cancer types (Fig. 2A). GSEA of their human homologues revealed relevant intratumoural pathways and processes (Fig. 2B-D). As expected ^21^, the upregulated transcripts were associated with interferon (IFN) cascades. Viral responses mediated by the 2′-5′ oligoadenylate synthase (OAS) proteins, that particularly sense cytosolic viral dsRNA ^22^, were remarkably enriched. The IFN-dependent ISG15 pathway, which closes the cell gates to DNA and RNA viruses by marking them through a post-translational modification ^23^, was also enriched. Biological processes were associated with response to viruses, cytokine production and signaling, and activation of innate immunity and inflammatory responses (Fig. 2C), while molecular functions highlighted double-stranded RNA binding and adenyltransferase activity (Fig. 2D). These findings suggest enhanced sensing and response to cytosolic dsRNA following CONV RT. We then reconstructed the physical and functional associations among the protein products of the 161 genes using STRING, to explore any protein-protein interactions. K-means clustering revealed a highly interconnected network of 64 proteins, with a prominent submodule (46 hubs) related to interferon signaling and response to viruses. This submodule entails the entire family of OAS proteins (OAS1, OAS2, OAS3, OASL), key components of the ISG15 machinery (ISG15 and USP18), STAT1 and interferon-stimulated genes (ISGs) which combat RNA and DNA viruses (Fig. 2E). A smaller submodule also includes proteins associated with striated muscle contraction (Fig. 2E). Although this may reflect stress-induced epigenetic alterations with lineage trans-differentiation, or myofibroblast differentiation from pericytes or fibroblasts of the TME, some recent studies also implicate sarcomeric proteins in radiosensitivity ^24^.

**Figure 2:**
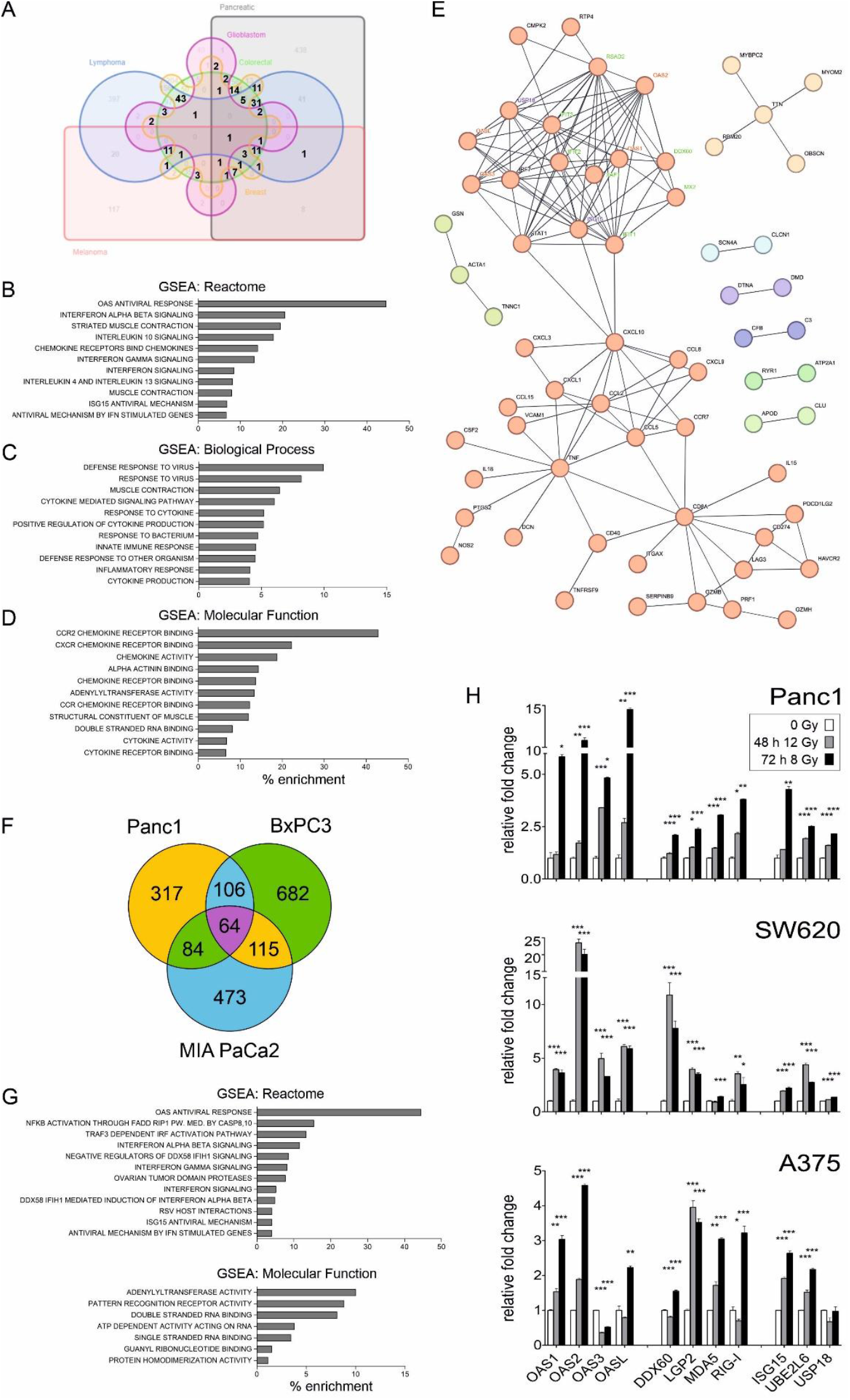
Activation of innate immune response of tumor cells to RNA viruses following CONV RT. (A) Venn diagram of CONV-RT upregulated gene transcripts per cancer type. Genes common across ≥ 3 cancer types are denoted with a number, with a total number of 141. (B-D) GSEA for reactome (B), biological process (C) and molecular function (D) of the 141 shared genes, (E) STRING network reconstruction of the interactions among the protein products of the 141 genes. The submodule of 46 red hubs are associated with IFN signaling and response to viruses, incliding OAS proteins, ISGylation proteins, and ISGs. The light-yellow submodule contains 5 proteins highly specific for striated muscle contraction. (F) Venn diagram of upregulated gene transcripts in Panc1, BxPC3, and MIA PaCa2 cell lines undergoing ICD 72 h after CONV-RT (3×8 Gy) reveals 64 shared genes (G) GSEA of reactome of the 64 genes, (H) qPCR for components of the OAS/RIG-I system and ISG15 pathway in irradiated Panc1 (pancreatic), SW620 (colorectal) and A375 (melanoma) cell lines 48h after 12 Gy and 72h after 8 Gy. RNA levels are presented relative to corresponding levels in non-irradiated cancer cell lines (0 Gys) after 72 h. Statistically significant differences with p value ≤ 0.05 (*), ≤ 0.01 (**) and ≤ 0.001 (***) are denoted.

Given that the abovementioned findings are inferred from the human homologues of RT-induced mouse genes, we validated their relevance in humans by using RNA-Seq data from irradiated human tumor cell monocultures known to exhibit robust cellular phenotypes of immunogenic cell death (ICD). CONV RT has been demonstrated to induce ICD in pancreatic cancer cell lines when administered as a 3×8 Gy regimen ^5^. By analyzing the corresponding transcriptomes (ArrayExpress, E-MTAB-13096), we found 64 significantly upregulated genes shared across the three irradiated pancreatic cancer cell lines 72h post-treatment (Fig 2E). These genes were enriched for the IFN-dependent viral innate immunity pathways OAS, RIG-I/MDA5, and ISG15 (Fig. 2F), as well as for molecular functions such as PRR, adenylase activity, and for RNA binding (Fig 2G). Furthermore, we irradiated human cell lines from three different cancer types (colon, pancreatic and melanoma) and monitored the transcriptional changes of key players of the abovementioned pathways (Fig. 2H), post 72h and 48h exposure to 1×8 Gy and 1×12 Gy, respectively. All candidates were actively transcribed within the tumour cell monocultures. For most candidates, induction was more robust at 8 Gy/72 h, implying that adequate time after irradiation may be required for the establishment of tumour transcriptional programs related to RT-induced immunogenic responses. Altogether, these findings suggest that immunogenic CONV RT doses facilicate intratumoural transcription of IFN-dependent pathways of viral immunity, with a particular preference for cytosolic sensors of RNA viruses.

### SFRT transactivates RNA viral sensors and NK cell responses earlier and more robustly than conventional radiotherapy

In the melanoma and the glioblastoma studies (Table 1, GSE268311 and GSE56113, respectively) mice were treated with SFRT or CONV RT, and tumour mRNA profiling expression was assessed via microarrays 2 days or 1 week (168-192h) post-irradiation. These datasets were analysed by utilizing our pipeline to detect conserved genes commonly responsive to SFRT in a time-dependent manner. An early common response pattern included the activation of members of the CCL and CXCL cytokine families, Toll-like receptors, and OAS proteins, all of which are involved in the recognition and/or response to pathogen-derived ligands (Supplementary Fig. 1A). At a later stage, 101 commonly SFRT-responsive DEGs were upregulated (Supplementary Fig. 1B), particularly enriched for DNAX-activating protein of 12kDa (DAP12) signaling, which promotes NK cell and DC cell cytotoxicity ^25^ (Supplementary Fig. 1C).

SFRT conferred immunogenic advantages and improved tumour control compared to CONV RT ^10-12^. To further delineate the SFRT-primed immunogenic signaling, we focused on the melanoma dataset, as the most recent and comprehensive study and incorporates more updated probeset annotations than the older glioblastoma studies. GSEA of the inferred human homologues revealed upregulation of a set of 76 transcripts at day 2, eight of which are commonly activated by SFRT and CONV, while the remaining are SFRT-specific. By day 7, 216 conserved transcripts were induced by SFRT and 227 by both SFRT and CONV. Notably, 70 of the 76 genes induced by SFRT at day 2 remained actively transcribed in SFRT-treated melanomas and only began to be expressed in the CONV-irradiated ones at day 7 (Fig. 3A). Comparison of the top-enriched pathways following SFRT indicated that the prominent earlier induced pathways are related to the OAS antiviral response, IL10 and interferon signaling, DAP12 signaling, and antigen processing cross-representation (Fig. 3B). Late SFRT responses are associated with the maintenance of IL10 activity and the induction of additional interleukins, such as IL2, IL6 and IL4/IL13 (Fig. 3B). In contrast, the CONV RT triggeredearly activation of ISG15 antiviral responses and interferon signaling, followed by late activation of the PD-1 and TCR signaling pathways at day 7 (Fig. 3B).

**Figure 3:**
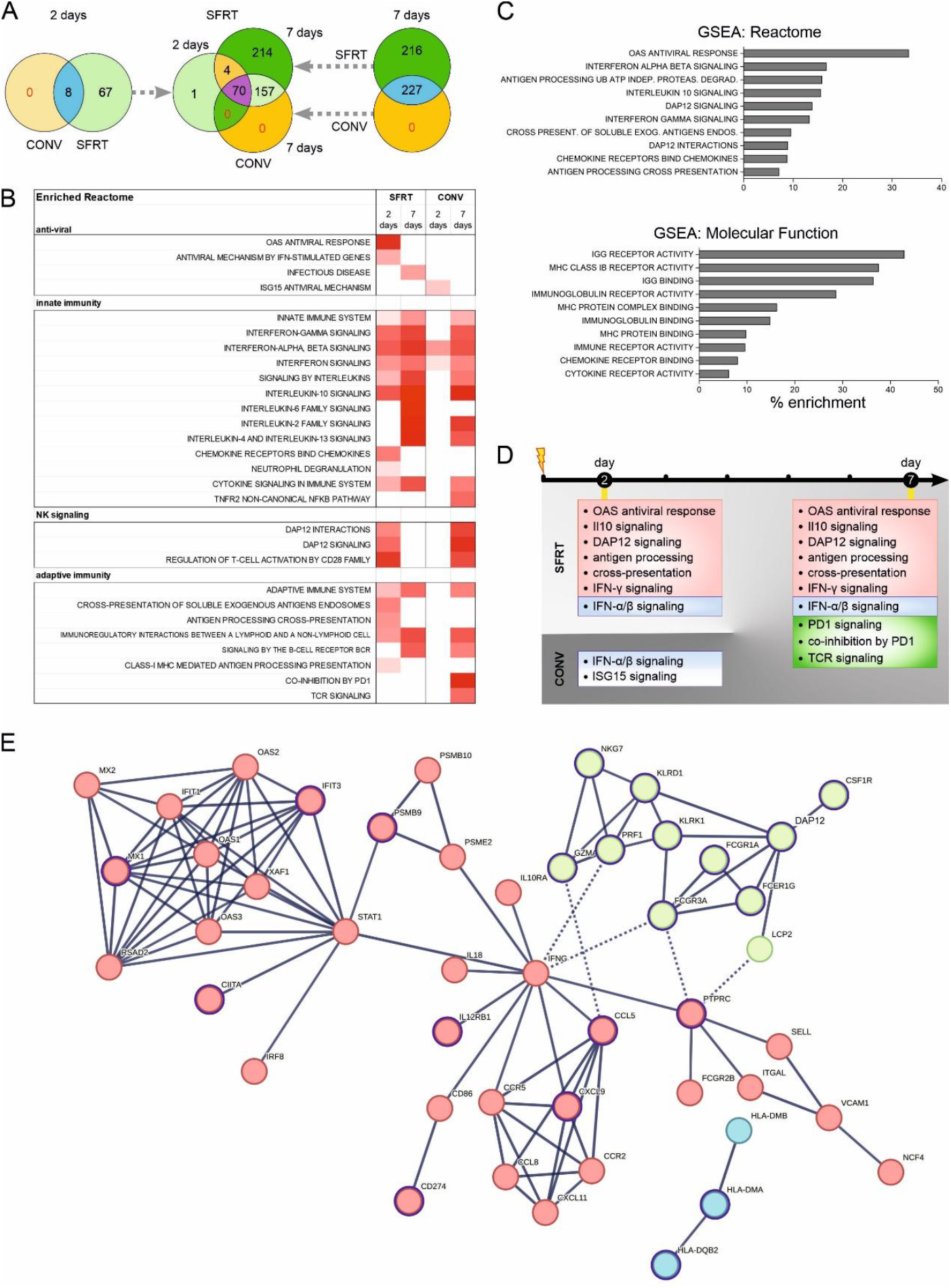
SFRT transactivates OAS antiviral and NK cell responses earlier and more robustly than CONV RT. (A) Venn diagrams of SFRT or CONV-RT upregulated gene transcripts in melanomas excised at day 2 (left) or day 7 (right) after irradiation treatments. Comparison of SFRT transcriptomes at days 2 and 7 with CONV RT transcriptomes at day 7 unveils 70 shared upregulated genes. (B) heatmaps of overrepresented reactomes 2 and 7 days after SFRT or CONV RT (C) heatmaps of overrepresented reactome processes of the 70 common genes (D) A timeline summary of activation of reactome processes by CONV RT vs SFRT (E) STRING analysis of the protein products of the 70 genes

The persistently upregulated set of 70 DEGs was enriched for pathways associated with the OAS antiviral response, ubiquitin-independent proteosomal degradation and DAP12 signaling (Fig. 3C), along with immunoglobin binding and activity (Fig. 3D). Their encoded proteins form a core network of 47 proteins, related to IFN alpha/beta signaling (red cluster, 33 hubs), NK cell-mediated toxicity (green cluster, 11 hubs) and peptide antigen assembly with the MHC II protein complex (blue cluster, 3 hubs). The components of the OAS viral response are clustered in a distinct submodule that connects to IFNγ via STATs. Several network components are associated with favorable prognosis in melanoma patients (Fig 3E). Finally, considering the coordinated enrichment of the OAS antiviral response and the DAP12 signaling, to which NK cells and myeloid DCs respond ^25^, we next examined whether OAS gene expression is associated with these cell types in patient melanomas. Analysis using TIMER2.0 ^26^ revealed that all OAS genes are positively correlated with NK (Fig. 4A) and DC (Fig. 4B) infiltration. Conversely, melanoma patients with OAS2 mutations exhibited decreased NK infiltration (Fig. 4C). Collectively, these results suggest that SFRT induces OAS-mediated RNA antiviral immunity, interferon responses and DAP12 signalling earlier and more robustly than CONV. The enhanced stimulation of these innate immune responses likely contributes to better tumour growth control following SFRT.

**Figure 4:**
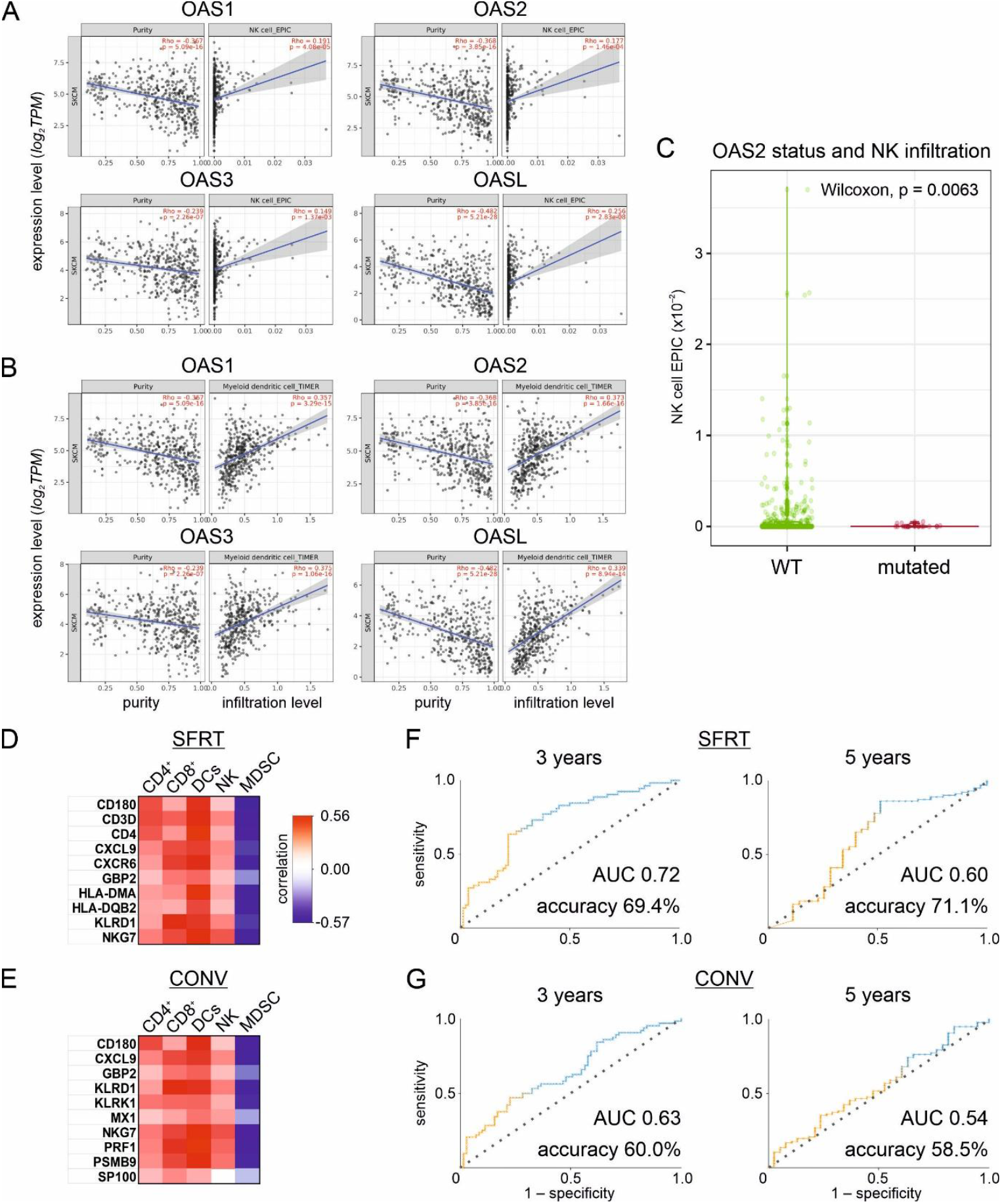
Gene expression signatures of response to SFRT are correlated with immune cell infiltration and better predictability of ≥ 3-year survival in melanoma patients. (A-B) Spearman correlation diagrams of RNA expression of OAS1, OAS2, OAS3, and OASL with NK cell (A) and DC (B) infiltration of melanomas from human patients. (C) association of OAS2 mutation and NK cell infiltration in human melanoma patients. (D-E) Heatmap of Spearman correlation values of the 10-gene signature for SFRT (D) or CONV (E) with CD4 T cells, CD8 T cells, DCs, NK cells, and MDSCs (F) ROC curves correlating the SFRT-induced immunogenic signature with survival of melanoma patients ≥ 3 years (left) and ≥ 5 years (right) after initial diagnosis, (G) Same as SFRT-induced immunogenic signature. Survival prediction accuracy is higher in the case of the SFRT signature, for both cut-offs.

### SFRT induces tumour signatures with higher predictability for longer patient survival than conventional RT

Unlike CONV RT, SFRT prolongs the survival of the irradiated mice ^10^. To investigate whether this association is clinically relevant, we sought to define characteristic signatures induced by each RT type and compare their ability to predict survival outcomes in human melanoma patients. Given the lack of microbeam studies in humans, we used survival data from the SKCM cohort of the TCGA, as a commonly referenced clinical setting. We hypothesized that if SFRT confers superior disease outcomes relative to CONV RT, then the SFRT-activated transcriptome should correlate more strongly with longer patient survival. In order to derive manageable signatures from the irradiated mouse transcriptomes, we prioritized the most significant upregulated transcripts by decreasing significance using XGBoost and Random Forest, identified their human homologues, and selected the top 10 homologues for each RT modality.

Using TIMER2.0, we assessed how the RNA expression of each gene in the CONV- or SFRT-specific signatures is associated with immune cell infiltration in the TME of melanomas in the SKCM TCGA cohort. The analysis included CD4+ and CD8+ T lymphocytes, DCs, and NKs which together shape an immunoreactive TME, as well as myeloid-derived suppressor cells (MDSCs), which promote immune evasion. Both signatures exhibited similar positive correlations with CD4+ T cells, CD8+ T cells, DCs and NKs, and negative correlations with MDSCs, implying an overall immune-hot TME, although the SFRT signature showed a stronger association with CD4+ T cells infiltration (Fig. 4D, E). We then performed multilayer perceptron modelling to estimate the predictive potential of the SFRT- and CONV RT-induced signatures for patient survivability at 1, 3 or 5 years after initial diagnosis. As indicated by the ROC curves (Fig. 4F, G), the SFRT-induced immunogenic signature performed better than the CONV-induced signature, especially for the 3-year period cut-off, with 69.4% accuracy (AUC: 0.72) versus 60% accuracy (AUC: 0.63). For the 5-year survival cut-off, the respective values were 71.1% (AUC: 0.60) and 58.5% (AUC: 0.54). Collectively, both characteristic signatures are associated with an immunoreactive TME, but the SFRT-induced one exhibited better predictive capacity for patient survivability.

### RT-induced intratumoural upregulation of viral RNA cytosolic sensors is accompanied by global changes in TE transcription

Considering that OAS and RIG-I, which are recurrently upregulated in irradiated cancer cells, require binding of cytosolic dsRNA in order to be activated ^27^, we searched for potential endogenous dsRNA molecules that could serve as their ligands. Retrotransposons, a heterogenous superclass of TEs ^28^, represent a important source for dsRNAs (Fig. 5A). TE transcripts can bind to RIG-I and induce an immune-hot phenotype, as observed in the oesophageal squamous cell carcinomas. LTRs were found to be the most abundant TE ligands of RIG-I ^6^. We hypothesized that this mechanism may extend to other cancer types, where irradiated cells undergoing ICD exhibit increased TE transcription, which could in turn act as the intratumoural stimuli of the overexpressed RNA sensors.

**Figure 5:**
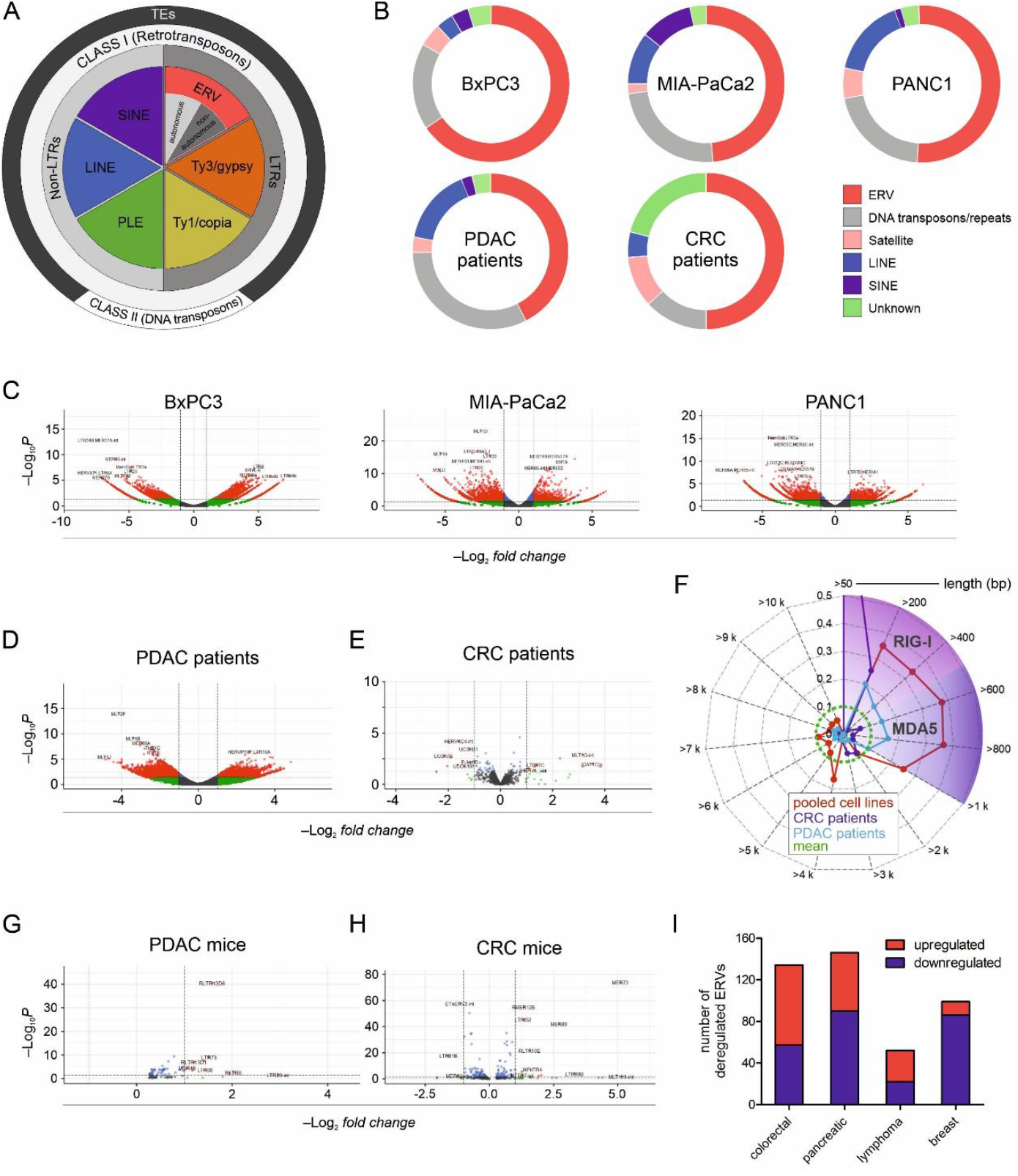
Global TE transcriptome changes in mouse and human tumours that exhibit RT-induced immunogenicity. (a) Classification of eucaryotic TEs. Class I (retrotransposons, including LTR/ERVs, LINEs and SINEs) require an RNA intermediate to transpose, and Class II (DNA transposons) move through a DNA intermediate. PLEs: Penelope-like elements, LINEs: Long interspersed nuclear elements; SINEs: Short interspersed nuclear elements (adapted and modified from ^28^ ), (B) Doughnut plots of the differentially expressed groups of TEs in irradiated vs. non-irradiated PDAC cell lines, PDAC patients and colorectal cancer patients. In all cases, the most prominent transcriptional changes are observed for the LTR family (mainly ERVs), followed by upregulation of DNA retroposons. (C-E) Volcano plots of deregulated expression of LTR/ERVs in PDAC cell lines (C), PDAC patients (D) and colorectal cancer patients (E), (F) Spider diagram of the length distribution of the upregulated autonomous LTR/ERV transcripts in the irradiated vs. non-irradiated pooled PDAC cell lines (red line), PDAC patients (blue line) and CRC patients (purple line) (G-H) Volcano plots of upregulated and downregulated LTR/ERV transcripts in irradiated vs non-irradiated mouse PDACs; red and green dots indicate up- and down-regulated genes, respectively, while gray and blue dots represent genes without statistically significant changes. (G) and colorectal tumors (H), (I) diagram of numbers of LTR/ERVS significantly deregulated in irradiated versus non-irradiated colorectal, pancreatic, lymphoma and breast mouse tumours.

TEs represent a substantial portion of the transcriptome but are generally ignored by the standard RNA-Seq pipelines, due to their repetitive nature and their incomplete mapping and annotation ^29^. For this reason, we preferably used TEtranscripts2, a specialized software that incorporates TE-associated reads in the differential expression analysis of human RNA-Seq data, discriminates between Class I (retrotransposons, including LTR/ERVs, LINEs and SINEs) and Class II TEs (DNA transposons), and has outperformed other similar tools ^29^. Analysis of RNA-Seq data from the pancreatic cancer cell lines ^5^, indicated global TE changes in irradiated tumour cells with the LTR family (mainly ERVs) exhibiting the most frequent changes, accounting for 49-65% of the differentially transcribed TEs. We evaluated the clinical relevance of this finding in surgically excised tumours from PDAC patients manifesting immune responses following RT compared to matched untreated patients ^30^, as well as in colorectal (CRC) tumours obtained from patients pre- and post-RT ^31^. The pattern of TE alterations in the irradiated tumours was consistent with that of the human PDAC cell monocultures (Fig. 5B). Collectively, these results indicate that immunogenic RT induces intratumoural upregulation of cytosolic RNA sensors, such as OASes and RIG-I-like receptors (RLRs), is accompanied by a massive accumulation of TE transcripts, predominately LTR/ERVs.

### Autonomous LTR/ERVs are re-expressed in mouse and human tumours that exhibit RT-induced immunogenicity

Most of the ERVs in the human genome are non-autonomous LTRs that do not encode proteins and primarily act as *cis* regulators. However, a small subset is autonomous, composed of LTRs flanking potential protein-coding sequences and approximating near full-length proviral sequences (Fig. 5A). The autonomous ERVs retain their ability to synthesize viral proteins ^29^, which can act as tumour-associated antigens eliciting T cell-mediated immunological recognition ^32^. Of note, anti-ERV antibodies have been shown to enhance cancer immunotherapy ^33^, while ERV activation has been associated with more favourable outcomes of clinical trial patients ^31^.

To investigate whether autonomous ERVs are transcribed within irradiated tumours, we utilized the ERVmap pipeline, which is highly suitable for the sensitive quantification of human ERVs from RNAseq data, due to its annotated database of 3.220 autonomous ERVs that approximate near full-length proviruses ^29^. We re-implemented the original Perl pipeline in Python and integrated up-to-date bioinformatics tools. Analysis of RNA-Seq data from irradiated human PDAC cell line monocultures (Fig. 5C), as well as PDAC (Fig. 5D) and CRC (Fig. 5E) patient tumours, revealed global deregulation of autonomous ERVs, with plenty of them being strongly upregulated. In PDACs, ERVs 400-600 bps and 600-800 bps long were most frequently upregulated, corresponding to RNA duplexes specifically recognized by the RIG-like receptors RIG-I (22-500 bps) and MDA5 (500-1000 bps) ^27^. CRCs also show an enhancement of ERVs at the range of 50-200 bps, a length correlated with peak activity of RIG-I (Fig. 5F). Hence, these findings suggest that RT induces ERVs with the potential to both activate RNA sensors via their transcripts and to be translated into proteins able of eliciting T-cell driven immune responses.

We further examined whether global ERVome changes are conserved between mouse and human irradiated tumours, by complementarily applying the Repenrich 2.0 pipeline ^34^ to the datasets in Table 1. Similar to the patient tumours, we observed genome-wide ERVome changes in the CONV-irradiated PDAC (Fig. 5G) and CRC (Fig. 5H) mouse tumours versus the non-irradiated controls. Similar changes were also observed in lymphoma and breast cancer (Fig 5I), indicating a cancer type-independent tendency of immunogenic RT to reactivate LTR/ERVs in mouse tumours. Unfortunately, similar analyses could not be performed for the RT-treated melanomas and glioblastomas, where the transcriptomic data were generated using microarray technology, which does not capture LTR/ERV transcripts.

### Enhanced abscopal effects are associated with common ERV activation and spreading of viral RNA sensing from irradiated to non-irradiated tumours

In the lymphoma study (Table 1, GSE281695), A20 cells were injected into both sides of the mice, and two irradiation regimens were tested separately, 1×8-Gy or 2×4-Gy. One of the developed tumours (IR) received irradiation, while the other tumour remained non-irradiated (NIR). Although both regimens exerted similar effects on the IR tumours, enhanced abscopal effects in the NIR tumours were reported only for the 2×4-Gy regiment, 7 days post-RT, but not for 1×8-Gy. Transcriptome analysis of the 2×4-Gy NIR tumour revealed activation of the same pathways as in the matched IR tumour, since 10 of the 12 top-enriched pathways in the IR (83.3%) were reproduced in the NIR tumour. Contrariwise, the 1×8-Gy sheme transactivated far fewer pathways in the NIR versus IR tumour (25%, 3 of the 12-top enriched in IR tumours) (Fig. 6A). Among the most enriched pathways shared between the 2×4 Gy IR and NIR tumours were the OAS-induced response to dsRNA viruses and the activation of C3 and C5 complement. Interferon pathways along with interleukin (IL0, IL4/IL13) and cytokine cascades were also recapitulated in the NIR tumour (Fig. 6A). Notably, the OAS-mediated sensing was not enriched in the 1×8-Gy NIR tumour transcriptome, despite the presence of interferon and interleukin pathways and the C3/C5 complement cascade. Moreover, half of the DEGs (116/242) detected in the 2×4 Gy IR tumours were also deregulated in the NIR tumours, compared to 17% (129/726) of the DEGs for the 1×8 Gy regimen (Fig. 6B). Therefore, the most efficient 2×4 Gy IR scheme achieved a higher reproducibility of the transcriptional changes and pathways between the IR and the NIR tumours. We further compared LTR/ERVome changes between the IR and NIR tumours. When the more efficient 2×4 Gy dose was applied, 62.9% of the LTR/ERVs activated in the IR tumours were also transcribed in the NIR tumours (Fig. 6C). In contrast, under 1×8 Gy, there were markedly fewer LTR/ERVs shared between the IR and NIR sites (16.6% versus 63%, respectively). Overall, the 2×4 Gy IR scheme promoted the dissemination of OAS-mediated sensing of dsRNA viruses along with transcriptional upregulation of ERVs common from IR to NIR sites.

**Figure 6:**
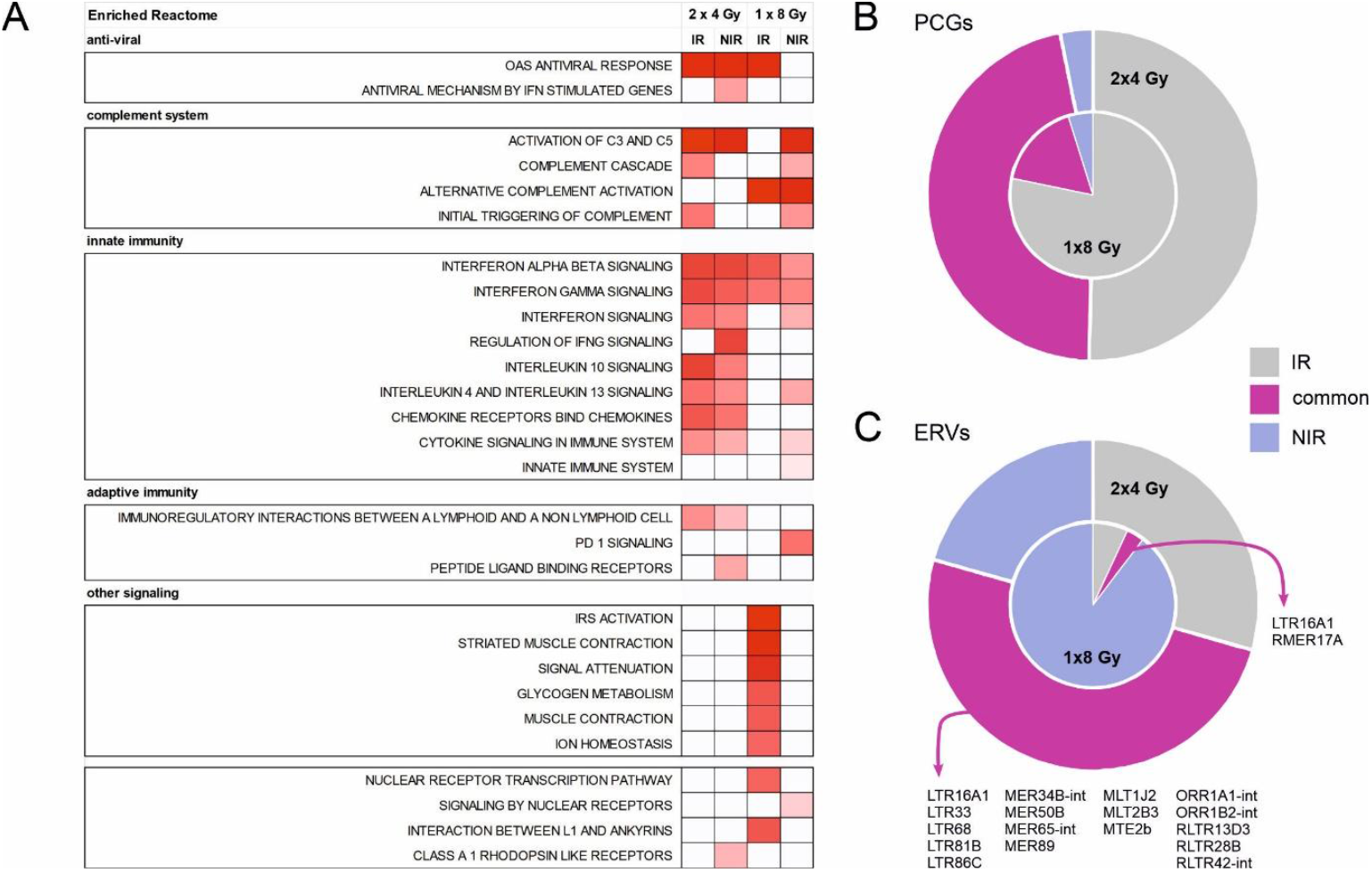
Enhanced abscopal effects are associated with transactivation of the same ERVs and viral RNA sensing in both IR and NIR sites. (A) Heatmap of overrepresented reactome processes based on the transcriptomes of lymphomas inside (IR) and outside (NIR) the irradiation field following treatment of mice with 2×4 Gy vs. 1×8 Gy. For the 2×4 Gy regimen, which elicited enhanced abscopal effects, OAS response is overrepresented both in IR and NIR sites. (B) circle diagrams of upregulated PCGs in the IR and NIR tumours treated with 2×4 Gy (outer circle) versus 1×8 Gy (inner circle). Purple segments represent the percentage of common PCGs (B) and LTR/ERVs (C) between IR and NIR tumors. In the case of 2×4 Gy, the purple segment is larger, representing greater overlap. (C) Same as (B) for upregulated LTR/ERVs.

## DISCUSSION

Herein, we show that CONV RT, at doses known to induce broad antitumor immune responses in vivo, transactivates IFN-I, OAS, and ISG15 pathways, along with IL-10 signaling. In the case of microbeams, an experimental form of SFRT, these pathways are activated earlier and more robustly along with NK cell responses, a fact that may be related to its superior performance relative to CONV RT. The SFRT-induced immunogenic signatures predict patient survival more accurately than those induced by CONV RT. We also identified conserved, global changes of TE Class I transcripts, particularly LTR/ERVs. In RT regimens that elicit abscopal effects, activation of the OAS dsRNA-sensing pathway and upregulation of the same LTR/ERV elements are observed in both IR and NIR sites. Parallel induction of LTR/ERVs and RNA viral sensors occurs inside the tumour cells.

RT is known to elicit intratumoural type I IFN responses ^21^. Members of the OAS family are conserved components of the IFN I pathway that recognize non-self dsRNA in a template length-dependent, but sequence-independent manner ^35^. In this way, they respond to viral infections while tolerating self dsRNA structures ^35^. Although transactivation of OAS genes is expected in this context, a breakthrough study suggests that they are not merely markers of IFN cascades but may also assert active roles in the establishment of an immunoreactive TME ^36^. OASes bind to non-self dsRNA in the cytoplasm to catalyze the production of 2-5A from ATP, which subsequently binds to and activates RNase L ^36^. The activated RNAse L cleaves RNA to smaller fragments that serve as RIG-I ligands, eventually amplifying IRF3/ IRF7-mediated transactivation and interferon response. Our observation that OASes and RLR receptors are co-elevated supports the existence of a possible dsRNA-triggered OAS/RIG-I/RNAse L axis of IFN activation. More intriguingly, 2-5A is a novel immunotransmitter that travels from donor to recipient cancer cells via gap junctions, thereby extending the reach of RNase L-mediated innate signaling ^36^. Therefore, transactivation of OAS genes may faciliate the paracrine transmission of innate immune signal across the irradiated tumour cells. This novel mechanism is particularly relevant to SFRT, where IR and NIR compartments of the tumour are in close spatial proximity. Based on our results, OAS-mediated signaling is persistently enhanced in SFRT-treated melanomas. A compelling hypothesis is that OAS-derived 2-5A is transported easier between alternating high- and low-dose regions, thereby propagating innate immune signaling more rapidy and ultimately accelarating the establishment of tumour immune responses.

Immunogenic responses to RT have traditionally been attributed to cytosolic dsDNA, which is sensed by the cGAS/STING pathway, whereas the co-elevated dsRNA fraction has received little attention ^37^. Nevertheless, RT increases IFNβ even in cGAS/STING-deficient cancer cells, indicating that cytosolic dsRNA accumulation may - alternatively or complementarily to dsDNA - activate IFN-I-mediated antitumour immunity, in certain contexts ^37^. Although the effect of the dsRNA-induced OAS-RNase L pathway on IFN production is relatively weak as compared to the most prominent cGAS/STING, it can be augmented selectively by the crosstalk between OASes and RIG-I ^38^, depletion of suppressors of the OAS-RNAseL interaction ^39^, and/or overexpression of the Sp1 transcription factor ^40^. Our findings are consistent with such upregulation of RNA-mediated IFN I response in irradiated tumours. This concept is further supported by the enrichment not only of RNA-specific PRRs, but also of other RNA-responsive pathways, such as ISG15 ^23^ and/or NK signaling ^41^.

The cytosolic accumulation of nucleic acids signals viral invasion and ignites cell-intrinsic antiviral pathways. In the absence of viral infection, these pathways can be activated by endogenous RNA originating, in part, from the re-suppression of otherwise silenced endogenous retrotransposons. The same PRRs detecting exogenous RNA viruses also sense retrotransposon RNA, establishing a ‘viral mimicry’ state that amplifies IFN response and eradicates cancer cells. Viral mimicry refers to the induction of antiviral responses by endogenous stimuli, such as retroelement-derived cytosolic nucleic acids, rather than by exogenous viral infection ^42^. Thus, TE transcripts repurpose the cellular defence machinery against RNA viruses to combat cancer ^7,43-47^. Since RT disrupts global epigenetic regulation ^48^, it could plausibly ‘awaken’ normally repressed repetitive elements ^7^, thereby increasing their transcription activity. These transcripts could, in turn, alarm for a virus infection at the tumour-bearing organ. In support to this notion, RT sparks ERV transcription and downstream activation of an innate antiviral MDA5/MAVS/IFN axis, thereby promoting antitumour immunity ^49^. Consistently, we found that following immunogenic RT doses, LTR/ERVs are co-upregulated with OASes and RLRs; this pattern is conserved in both human and mouse tumours. Although the binding specificities of these PRRs for distinct LTR classes have not yet been characterized, the LTR/ERVs transcripts may serve as cues of the upregulated RNA sensors and have been shown to bind to RIG-I ^6^.

ERVs are remnants of ancient retroviruses embedded in the so-called ‘junk DNA’ regions of the human genome. They co-evolved with humans and established a complex relationship with the immune system. During tumorigenesiss, ERVs lose their strictly controlled spatiotemporal silencing, and their promoter/enhancer activity can be hijacked by oncogenes. Nevertheless, this out-of-context activation can backfire for the tumour, because the products of re-expressed ERVs can be recognized as viral threats. At the RNA level, ERV-derived transcripts activate IFN-regulated innate immune response, whilst at the protein level, ERV peptides may act as neoantigens, triggering B-cell mediated responses ^50^. By awakening ERVs, RT may generate a tumour-intrinsic reservoir of antigenic molecules that recruit innate and adaptive immunity against the tumour. ERVs-generated antigens drive humoral B cell responses from tertiary lymphoid structures (TLS), which are increasingly recognized as favorable biomarkers for immunotherapy responses. It would be of interest to investigate whether RT-induced ERV expression primarily launches a humoral B cell response, establishing or maturing existing TLS structures that complement and boost T-cell driven immunotherapy responses ^50^.

Conventional homogenous RT delivers the maximum tolerated dose but can also initiate adverse events. In contrast, SFRT leverages the distinct biological effects of varying doses to optimize the simultaneous activation of multiple immunogenic effects in a single TME, preserve antitumour immunity, and at the same time mitigate toxicity in healthy tissues ^14^. The superiority of SFRT relies on the spatial heterogeneity of immune features. For example, administration of heterogeneous RT on tumour-bearing mice by high-dose-rate brachytherapy demonstrated superior in situ vaccine effects, especially in combination with immunotherapy ^51^. Even more intriguingly, when one half of the tumour receives high and the other half low dose, bilateral crosstalk is established between the heterogeneously irradiated regions, boosting immune responses ^14^. In the case of minibeam or microbeam irradiation, whereby the repetitive pattern of peaks and valleys generates an array of alternating high-and low-dose regions within the same tumour, the number of such favorable interactions is theoretically maximized, in proportion to the high/low-dose interfaces. In melanoma, SFRT does not induce transcriptional programs that are markedly distinct from those of CONV RT; rather, it triggers IFN I-associated antiviral pathways early and more robustly, with particularly accelerated activation of OASes. It would be of interest to investigate whether this temporal feature contributes causally to the superior efficacy of SFRT.

Our study represents the first systematic approach to correlate organismal-level phenotypes of broad antitumour immunity with tumour transcriptional reprogramming upon exposure to homogenous or spatially heterogenous RT, offering insights into the underlying mechanisms. We demonstrated that RT-induced immunogenicity is associated with tumour cell-intrinsic activation of IFN-dependent antiviral pathways, that respond mainly to RNA viruses. These findings provide a framework for the rational design of future functional studies investigating the mechanisms of abscopal effects. Viral mimicry via RT-induced ERV de-repression emerges as an appealing strategy to boost antitumour responses, but several questions remain open. First, it is unidentified which TE(s) bind to which innate RNA sensor(s). Second, the conditions under which awaking of ERVs turns beneficial for the host, especially for promoting abscopal effects, need thorough characterization, since ERVs could also act as double-edged swords favoring autoimmunity ^42^, aging ^52^, neurodegenerative disorders ^53^ or (upon chronic expression) evolution of tumour immune evasion ^54^. Third, the systemic signals released by irradiated tumour cells to recapitulate ERV and OAS-mediated responses at distant sites remain to be fully characterized. Furthermore, optimization of SFRT parameteres such as geometry and size of the high-dose regions is essential to ensure maximal antitumour immunity while sparing healthy tissue ^9^. Single-cell and spatial omics can capture the heterogeneity and architecture of the irradiated tumours. ERVome profiling and RNA-protein interaction analyses can uncover non-coding RNA-mediated pathways. Application of these interdisciplinary methodologies provides a fertile ground for elucidating the key underlying mechanisms and advancing personalized patient management.

## MATERIALS AND METHODS

### Mouse studies for phenotype-transcriptome correlations and identification of human orthologues

We queried the NCBI Gene Expression Omnibus (GEO) datasets using the following combinations of keywords: (“mouse” and “in vivo tumour immune response” and “irradiation”), (“mouse” and “carbon ion” and “tumour immune response”), (“mouse model” and “irradiation” and “immunogenicity”), (“mouse” and “irradiation” and “antitumour immunity”), (“irradiation” and “antitumour immunity”), (“murine”, “irradiation”, “anti-tumour immunity”), (“spatial fractionated irradiation”). We extracted all studies where mice exhibited broad antitumour responses to RT, either alone or in combination with immunotherapy, and for which bulk RNA sequencing (RNA-seq) data from both irradiate d and non-irradiated (untreated) tumours are provided (last accession date: March 20, 2025). The differentially expressed genes (DEGs) in irradiated tumours compared to non-irradiated controls were detected through RNAseq analysis. The human homologues of the significantly upregulated mouse genes (log2FC >1.5, p value < 0.05) were identified as previously described ^55,56^.

### Bulk RNA sequencing analysis

The SRA files were downloaded from NCBI GEO and converted into FASTQ using the Sequence Read Archive (SRA) Toolkit v.3.0.0 (available at https://github.com/ncbi/sra-tools) with the *fasterq-dump* utility. Adapters and low-quality bases were removed from the raw RNA-Seq reads using Trimmomatic and the filtered reads were then aligned to either the mouse (mm10) or human genome (hg38) using STAR v2.7.1.1. Alignment files were sorted using Samtools and transcript abundance was quantified with StringTie ^57^. Samples were clustered based on their expression profiles with Principal Component Analysis (PCA) in an R environment. Differential expression analysis was performed using the DESeq (v2 1.12.3) ^58^ package in R.

### TE identification from RNA-Seq data

Transcript-level TE quantification was performed using TEtranscripts (v2.2.3) ^59^. A dedicated Conda virtual environment was created using Miniconda3, incorporating Python (v3.8), pysam (v0.16.0.1), and R (v4.3.0). The FASTQ files corresponding to the GSE207717, GSE185311, and GSE179351 datasets were obtained from NCBI GEO, and, for each dataset, the sorted BAM files generated as intermediate outputs from the ERVMAP (v1.1) pipeline were used as input for this analysis ^29^. TE annotations were derived from the GRCh38_Ensembl_rmsk_TE.gtf file and its associated hg38_rmsk_TE.gtf.ind index file, both curated and publicly released by Oliver Tam (NYU Langone Health; released on May 7, 2024).

TEtranscripts was executed in multi-mapping mode to ensure quantification of reads mapping to both genes and repetitive elements. Differential TE expression analysis between the RT-treated and untreated (control) samples was conducted using DESeq2 version 1.12.3 ^58^. For the GSE207717 dataset, three biological comparison groups (BxPC3, MIA-PaCa2, PANC1) were defined, each consisting of three treatment and three control samples. The GSE185311 dataset included 9 SBRT-treated and 13 untreated samples, while the GSE179351 dataset comprised 6 treatment and 13 control CRC samples.

The transcript-level quantification was complemented with TElocal (v1.1.1) for locus-specific expression analysis of transposable elements ^59^. The previously established Conda environment was reused, with Python (v3.8) and pysam (v0.16.0.1), thus complying with TElocal’s software requirements. Input data consisted of the sorted BAM alignment files generated by ERVmap (v1.1) ^29^. Locus-level annotations were provided by the GRCh38_Ensembl_rmsk_TE.gtf and the corresponding GRCh38_Ensembl_rmsk_TE.gtf.locInd locus index file (courtesy of Oliver Tam). TElocal was executed in multi-mapping mode, processing each sample iteratively using a SLURM job submission script. Dynamic output directories and project-specific naming conventions were created for each sample to maintain data integrity and traceability. HERVd ^60^, a database of human endogenous retroviruses, was utilized for assigning each element to its corresponding class (i.e. ERV, LINE, Retroposon, Satellite etc.).

### Identification of human autonomous LTRs from transcriptomic data

To identify autonomous LTR elements, we modified a previously published Perl-scripted pipeline^29^. Specifically, we re-implemented the original Perl script in Python by integating updated bioinformatics tools for adapter trimming, sequence alignment, multithreading, error handling and dependency management. This reimplementation improved efficiency, readability, and reproducibility while maintaining full compatibility for ERV detection and quantification using paired-end RNA sequencing data. The updated Python pipeline is available at: https://github.com/IBG-ComputationalSystemsBiology/erv-mapping-pipeline.

### Quantification of repetitive elements from mouse RNA-Seq data

RepEnrich2 was installed according to the guidelines available at https://github.com/nerettilab/RepEnrich2. The software and its dependencies were installed in a cluster computing environment, including Python 2 v2.7.9 (Foundation), Bowtie2 v2.3.5.1, Bedtools v2.29.2, Samtools v1.5, and BioPython v1.76. Repetitive element annotations were obtained from RepeatMasker ^61^ and were downloaded in BED file format compatible with the *Mus musculus* genome (mm9). The annotation setup was executed using the script *RepEnrich2_setup*.*py*, provided within the RepEnrich2 modules. The RNA-Seq reads were aligned to the reference genome using Bowtie2. Uniquely and multi-mapping reads were filtered using Samtools and the *RepEnrich2_subset*.*py* script. This script was modified by adding the ‘Other’ class for reads not matching any LTR in the annotation file. Reads were then assigned to repeat families to quantify their expression levels, and the resulting count tables were compiled in a single Excel file containing all biological and technical replicates. After filtering out the ‘Other’ category, the identified LTRs were retrieved from the annotation file. Differential expression analysis was subsequently performed using DESeq2 v1.12.3 ^58^ in an R environment.

### Machine learning for identifying characteristic signatures of response to CONV or SFRT

Random Forest and XGBoost were applied to rank the most significant DEGs from each mouse study and cancer type, for each delivery mode, i.e. CONV or SFRT. All analyses were conducted in R v4.4.0. For Random Forest, a model was built using the randomForest (v4.7-1.2) package. The target (dependent) variable was the signature score, while the predictors (independent variables) included the weighted logFC (0.4) and weighted time (0.6) to balance expression change and exposure time. The model was trained with 500 trees (ntree = 500), and variable importance was assessed using the percentage increase in mean squared error (%IncMSE), reflecting the contribution of each variable to the predictive performance. Importance scores were calculated for each gene and integrated with the original weighted values to compute a combined score capturing both the expression magnitude and time-dependent effects. The top-ranked genes based on the combined score were considered as the most significant. For XGBoost, a model was built using the xgboost (v1.7.10.1) package. A synthetic target variable was defined as the sum of the weighted logFC and time. The resulting score was used to build a regression model trained on the weighted features, with the objective function set to squared error (reg:squarederror). The model was trained for 100 boosting rounds with a learning rate of 0.1 and a maximum tree depth of 6. Feature importance, calculated using the *xgb*.*importance* function, was used to rank genes as candidate signature genes.

### Survival analyses, GSEA, protein Interaction Networks and tumour infiltration analyses

Cox regression analysis of The Cancer Genome Atlas (TCGA) transcriptomics, Gene Set Enrichment Analysis (GSEA) and protein interaction network construction by STRING have been described previously ^55,56^. For GSEA, the Gene Ontology (GO) terms were plotted according to their enrichment percentage. For tumour infiltration analysis, the Spearman’s ρ correlation coefficients between the expression levels of each gene and the estimated infiltrate levels of selected immune cell types in the TCGA cohorts of melanoma patients (primary and metastatic) were acquired through the Tumour Immune Estimation Resource 2.0 [TIMER2.0] ^26^.

### Multilayer perceptron modelling for correlation of gene signatures with patient survivability

Using the characteristic top 10 human gene homologues of CONV RT or SFRT, we modelled the expression of the ranked genes in the tumours from the SKMC TCGA cohort to predict cancer patient survival at > 1, 3 or 5 years after diagnosis. A classification model for each survival period was built using a multilayer perceptron (MLP), i.e. a neural network composed of an input layer, hidden layers, and an output layer, whereby each layer contains a set of neurons. For each survival period, 2 hidden layers with 50 nodes per layer were used. The input features were normalized to a range of 0 to 1 using the machine learning software platform *weka 3*.*8*.*6* (University of Waikato, New Zealand). A random test set comprising 30% of the data was selected for evalutaing the performance of the model. To avoid overfitting, the training data were expanded to triple their original size by adding Gaussian noise with mean=0 and standard deviation=0.1; the test data remain unchanged. In the study group, the survival rates in the patient cohort for 1, 3 and 5 years were 87.46%, 50.78% and 36.99%, respectively. The network was trained for 500 epochs with a learning rate of 0.3. The model performance was evaluated using the area under the curve (AUC) of the Receiver Operating Characteristic (ROC) curve by plotting the true positive rate (TPR) and the false positive rate (FPR) across different thresholds.

### Cell cultures and irradiation treatments

Panc 1 cells were cultured in RPMI 1640, while SW620 and A375 cells in DMEM, supplemented with 10% Fetal Bovine Serum and 1% penicillin-streptomycin (Thermo Fisher Scientific). For the X-ray treatments, SW620 (10^6^ cells), Panc1 (500,000 cells) and A375 (250,000 cells) were plated in 10 cm dishes. After 24h, when the cells reached a confluency of 60-70%, they received either 8 Gy or 12 Gy radiation dose in a MultiRad225/26 irradiator (Faxitron Biotics), using the following settings: 200 kV X-rays; 17.8 mA; 0.5 mm Cu-filter; 2.151 Gy/min, 37 cm distance from the source, and 20 cm field size. The irradiated cells were then incubated at 37°C and 5% CO_2_.

### RNA isolation, reverse transcription and real-time PCR

After 48h (for the 12 Gy treatments) or 72h (for 8 Gy treatments) of irradiation, cells were harvested. Total RNA was extracted using the RNeasy kit (Qiagen) according to the manufacturer’s instructions. 1 µg total RNA per sample was reverse-transcribed using the RT2 First Strand Kit (Qiagen). For real-time PCR, the cDNAs (20ng) were mixed in a 20 µL reaction consisting of nuclease-free water, 20 pmol of each primer pair, and 7.5µL of Power SYBR Green PCR Master Mix (Thermo Fisher Scientific). Amplification with the appropriate primers (Supplementary Table) was performed in a StepOnePlus Real-time PCR system (Thermo Fisher Scientific) using the following conditions: 95 °C for 10 min (1 cycle); 95 °C for 15 sec, primer-specific annealing temperature for 30 sec, 72°C for 45sec (40 cycles). Statistical analyses were conducted using a two-sided Student’s t-test.

### Code availability

The Python script is available at: https://github.com/IBG-ComputationalSystemsBiology/erv-mapping-pipeline

### Data availability

The transcriptomics data analyzed in these study are publicly available in the Gene Expression Omnibus (GEO) repository under the following accession numbers: GSE56113, GSE61208, GSE168016, GSE179351, GSE185311, GSE207717, GSE209765, GSE255242, GSE268311, GSE281695, and in ArrayExpress repository under the number E-MTAB-13096.

## Author Contributions

**Stella Logotheti:** Conceptualization, Data curation, Formal analysis, Funding acquisition, Investigation, Methodology, Visualization, Writing – original draft, Writing – review and editing, **Elif Yildiz**: Software, Investigation; **Sarah Hasan:** Software, Investigation; **Elpida Theodoridou:** Validation **Stefan Kuhn:** Resources, Software, Methodology **Thorsten Stiewe:** Writing – review & editing **Stephan Marquardt:** Methodology, Data curation, Visualization, **Athanasia Pavlopoulou:** Data curation, Formal analysis, Investigation, Methodology, Writing – original draft, Writing – review and editing, **Joao Seco:** Conceptualization, Funding Acquisition, Supervision. All authors approved the final version of the manuscript prior to submission

## Acknowledgement

Figure 1A was created using BioRender (biorender.com).

## Funding

TS acknowledges funding support from Deutsche Forschungsgemeinschaft (STI 182/15-1), Deutsche Krebshilfe (70116475), and Wilhelm-Sander Stiftung (2022.129.1). SL and SK acknowledge COST Action CA21169 DYNALIFE. SL and JS acknowledge support from Deutsche Forschungsgemeinschaft (LO 3128/4-1, Projektnummer: 561811975) and DKFZ Collaborative program 2025.

## Conflict of Interest

The authors declare that they have no conflict of interest

## Additional information

The following are the Supplementary data to this article:

**Supplementary Table 1:** Sequences for qPCR primers

**Supplementary Figure 1: Upregulated genes and pathways in SFRT-treated tumors**. (A-B) Venn diagrams of human orthologues of the genes that are transcriptionally elevated in melanomas and glioblastomas two (A) or seven days (B) after receiving SFRT (microbeam). Members of the same protein families are highlighted in red lettering (C) GSEA for the reactome of the 101 shared upregulated genes between SFRT-treated melanomas and glioblastomas seven days after therapy.

## Notes

### Competing Interest Statement

The authors have declared no competing interest.

