## Supplementary Table 1 for "In situ vaccine effect of radiotherapy is associated with intratumoural ERV transcription and RNA virus sensing"

Supplementary Table 1: qPCR primer sequences

| **Primer name** | **Sequence** |
| --- | --- |
| **OAS1 Forward** | GAAGCTGCCACCTCAGTATG |
| **OAS1 Reverse** | TTCCAAGACCGTCCGAAATC |
| **OAS2 Forward** | TGGCTCCTATGGACGGAAA |
| **OAS2 Reverse** | GATGTCACGTTGGCTTCTCT |
| **OAS3 Forward** | CCCAAGCCACAAGTCTACTC |
| **OAS3 Reverse** | TGGGCGAATGTTCACAAAGT |
| **OASL Forward** | GCTGAAGGTAGTCAAGGTGG |
| **OASL Reverse** | CTCCTGGAAGCTGTGGAAAC |
| **ISG15 Forward** | GCAGATCACCCAGAAGATCG |
| **ISG15 Reverse** | GGCCCTTGTTATTCCTCACC |
| **UBE2L6 Forward** | CCCTCAATGTGCTGGTGAAT |
| **UBE2L6 Reverse** | GGTGAACTCTTCGGCATTCT |
| **USP18 Forward** | TGGGAAACAGGTCTTGAAGC |
| **USP18 Reverse** | CAGGGAGTGGCAGATCTTTC |
| **RIG-1 Forward** | AGAGCAAGAGGTAGCAAGTG |
| **RIG-1 Reverse** | ATACTGCTTCGTCCCATGTC |
| **MDA5 Forward** | GAGGAATCAGCACGAGGAAT |
| **MDA5 Reverse** | ATCAGATGGTGGGCTTTGAC |
| **DDX60 Forward** | TGTTTGTCTGTCTGGGAACT |
| **DDX60 Reverse** | AAGCGACATTTTCCTTCCTC |
| **LPG2 Forward** | GCCTTGCAAACAGTACAACC |
| **LPG2 Reverse** | AATTTCCGGCTCAACTCAGG |
| **RPL0 Forward** | GGTCATCCAGCAGGTGTCC |
| **RPL0 Reverse** | CAGACACTGGCAACATTGC |
| **GAPDH Forward** | GCATCCTGGGCTACACTGAG |
| **GAPDH Reverse** | AAAGTGGTCGTTGAGGCA |
