## Supplementary figures and images for "In situ vaccine effect of radiotherapy is associated with intratumoural ERV transcription and RNA virus sensing"

### Supplementary Figure 1

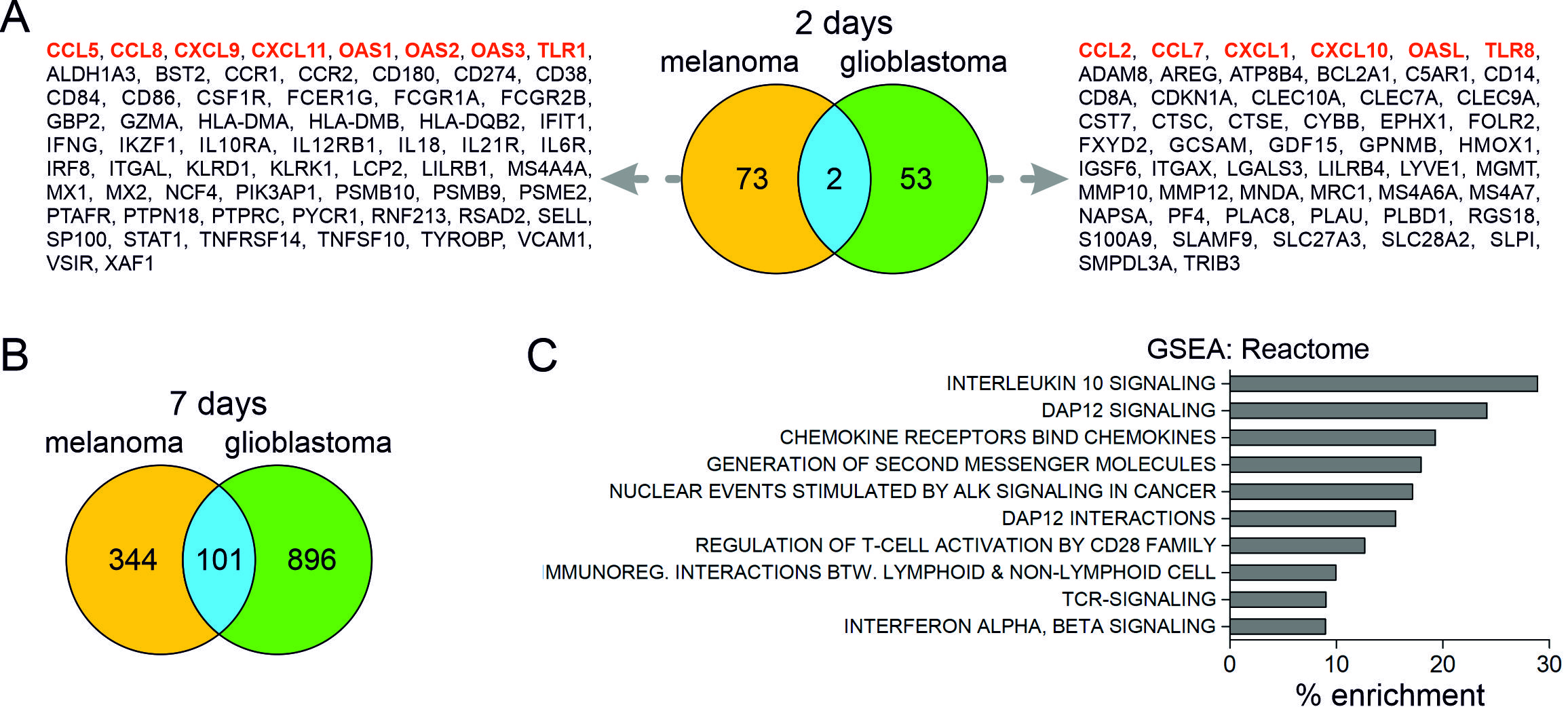
